# Toward pharmacologic therapy for glioblastoma: Characterization of the very long-chain acyl-CoA synthetase 3 (ACSVL3) inhibitor Grassofermata

**DOI:** 10.64898/2026.07.07.736493

**Authors:** Emily M. Clay, Xiaohai Shi, Elizabeth A. Kolar, Yanqiu Liu, Bachchu Lal, Paul A. Watkins

**Author notes:** Please address correspondence to: Paul A. Watkins MD, PhD Kennedy Krieger Institute 707 N. Broadway Baltimore, MD 21205 USA 1-443-923-2754.

## Abstract

Malignant brain tumors are among the most aggressive and difficult to treat human cancers. Glioblastomas (World Health Organization grade IV gliomas) are particularly lethal and refractory to treatment. Few drugs exist that are even somewhat effective. Our investigation of the physiologic role of fatty acid (FA) activating enzymes (acyl-CoA synthetase; ACS) identified an ACS that was widely expressed in gliomas but not in normal glial cells. Depletion of this enzyme, ACSVL3 (very long-chain ACS3), by knockdown or knockout decreased the malignant behavior of several glioma cell models including U87MG and Mayo-22 cells both in culture and when grown as xenografts. Hypothesizing that ACSVL3 is a potential therapeutic target in glioma, we conducted a search for inhibitors of this enzyme and found that CB5 (grassofermata) was a promising candidate. Treating U87MG glioma cells with CB5 slowed growth in monolayer culture; the growth rate was similar to that seen in cells in which ACSVL3 was either knocked down or knocked out. CB5 inhibited growth in a dose-dependent manner over a narrow range, and concentrations above 10 µM were toxic. Treatment at the lower dose of 3 µM inhibited growth of U87MG cells but was reversible, suggesting that this dose was not toxic. CB5-treated U87MG cells exhibited an altered morphology with a larger size and longer projections. In contrast, normal human fibroblasts treated with 10 µM CB5, a concentration that was toxic to U87MG cells, showed no effect on either growth rate or morphology. Treating U87MG cells with 3 µM CB5 induced differentiation as shown by increased expression of the astrocyte-specific marker glial fibrillary acidic protein (GFAP). In contrast, GFAP levels remained low in ACSVL3 knockdown cells. CB5-treated U87MG cells were less invasive, and thus less malignant, than either untreated cells or ACSVL3 knockout cells when assessed by a scratch wound healing assay. Acute treatment of U87MG cells with 3 µM CB5 decreased the ability of these cells to degrade FA of differing chain lengths from 16-24 carbons by β-oxidation, suggesting that decreased ACS enzyme activity contributes at least in part to the drug’s mechanism of action. NOD/SCID mice receiving up to 32 mg/kg/day CB5 by intraperitoneal injection showed no obvious side effects, suggesting that the drug was well-tolerated. Xenografts induced by subcutaneous injection of U87MG cells in the flanks of NOD/SCID mice were allowed to grow for 8 days after which half of the mice were treated with 2 mg/kg/day CB5. After 7 days of treatment, xenograft growth slowed in the treated mice and by 12 days tumor size had begun to decrease, suggesting therapeutic efficacy. When a similar study was done using xenografts induced by subcutaneous injection of Mayo-22 cells, which are maintained as subcutaneous tumors in mice rather than in cell culture, the effect of CB5 on tumor growth or weight at sacrifice was not statistically significant. The results of these studies suggest that CB5 may have therapeutic value in malignant glioma. Additional studies using other glioma models and other drugs chemically related to CB5 seem warranted.

## INTRODUCTION

Altered lipid metabolism is among the many changes observed in cancerous cells [Bian et al., 2021; Cheng et al., 2018; Snaebjornsson et al., 2020]. We previously reported that the lipid metabolism enzyme very long-chain acyl-CoA synthetase 3 (ACSVL3; also called FATP3) is overproduced in several cancers, including malignant gliomas and lung tumors [Pei et al., 2009; Pei et al., 2013]. A recent study suggested that ACSVL3 levels are also elevated in pre-leukemic and leukemia cells [Liu et al., 2024). Gliomas, and particularly World Health Organization grade 4 gliomas (glioblastoma; GBM), are aggressive cancers that are resistant to most treatment modalities. The standard of care is administration of the chemotherapeutic drug Temozolomide along with surgery and radiation therapy [Stupp et al., 2005]. Even with excellent clinical care, most GBM patients live only 12-18 months following diagnosis, and 5-year survival is only around 7% [Louis et al., 2016; Ellison, 2016; Ostrom et al., 2023]. It is clear that better treatment options, including better drugs, are needed to treat glioma patients and improve survival rates.

Targeting fatty acid metabolism is a promising therapeutic option in gliomas [Miska and Chandel, 2023]. To determine the relationship between ACSVL3 and glioma, we investigated its cellular and biochemical properties. In addition to tumor tissue, ACSVL3 levels were found to be higher in human glioma-derived cells, including U87MG and Mayo-22 cell lines [Pei et al., 2009]. Both of these cell lines exhibit malignant behavior in culture, and are tumorigenic in mice. Decreasing the expression of ACSVL3 in these cell lines using either knockdown or knockout (KO) strategies resulted in a more “normal” growth phenotype and a lower tumorigenic potential [Pei et al., 2009; Kolar et al., 2021]. This suggested that an inhibitor of ACSVL3 might be a candidate therapeutic agent in glioma.

Enzymatically, ACSVL3 is an acyl-CoA synthetase (ACS). ACSs catalyze the thioesterification of coenzyme A (CoA) to a fatty acid (FA), thereby activating the FA for its participation in essentially all further downstream metabolic processes. FA-CoA, and not unesterified FA, are substrates for catabolic (e.g. FA β-oxidation) and anabolic (e.g. complex lipid synthesis) pathways [Watkins 1997]. ACSs are therefore key players in FA metabolism. In part because of the diversity of chain lengths of FA in nature, subfamilies of ACSs that prefer FA substrates of different chain length have evolved. The very long-chain (ACSVL) family includes six structurally related enzymes (ACSVL1-6; gene names SLC27A1-6) that are capable of activating FA containing ≥20 carbon atoms [Watkins et al., 2007]. We reported that ACSVL3 preferentially activated saturated FAs containing 18-22 carbons [Kolar et al., 2021]. While lowering cellular ACSVL3 levels in KO (U87-KO) cells did not appreciably affect rates of FA synthesis or oxidation, it did significantly alter sphingolipid metabolism and related signaling pathways [Kolar et al., 2021].

Because no highly specific inhibitor of ACSVL3 is currently known, we used a two-pronged approach to identify potential ACSVL3 inhibitors. Our first approach was to test compounds previously shown to inhibit the closely related enzyme, ACSVL1 (also called FATP2) [Li et al., 2008; Sandoval et al., 2010]. Black and coworkers identified several inhibitors of ACSVL1, and we tested them for their ability to inhibit ACSVL3 [Clay et al., 2025]. Secondly, we developed a high-throughput screening assay for identifying candidate ACSVL3 inhibitors based on the strategy used in the Black laboratory to identify compounds that inhibit ACSVL1 [Kolar et al., 2021]. One ACSVL1 inhibitor that stood out as a good candidate ACSVL3 inhibitor was CB5 [Clay et al., 2025], also known as grassofermata [Black et al. 2016; Saini et al., 2015], Here, we report the results of both in vitro and in vivo studies of the therapeutic potential of CB5 to modulate the malignant behavior of cultured U87MG cells and the growth of subcutaneous xenografts arising from implanted U87MG cells.

## MATERIALS AND METHODS

### Materials and general methods

Cell culture reagents were from CellGro Technologies except for fetal bovine serum (FBS) which was from Biosource International. All [1-^14^C]fatty acids, including palmitic (C16:0), stearic C18:0, oleic (C18:1), behenic (C22:0 and lignoceric (C24:0), were from Moravek, Inc. Unlabeled fatty acids were from Millipore Sigma. DMSO was from Thermo Fisher Scientific. Cell authentication for U87MG cells was done within a year of study by the Johns Hopkins Genetics Resources Core Facility by short tandem repeat analysis using the PowerPlex 16 HS kit (Promega). Protein was measured by the method of Lowry et al. [1951] unless otherwise indicated. CB5 and CB16.2 were provided by Drs. Paul Black and Concetta DiRusso (Department of Biochemistry, Univ. of Nebraska, Lincoln NE). All were prepared as 1000x stock solutions in DMSO and added to cultures and assays to the final concentration indicated in each experiment. The final DMSO concentration was 0.1%.

### Cell culture

U87MG cells were obtained from American Type Culture Collection. The ACSVL3-knockout (U87-KO) cell line was produced with the CompoZr Knockout Zinc Finger Nuclease kit (Millipore Sigma) and characterized as described previously [Kolar et al., 2021]. Both lines were cultured in Dulbecco’s Modified Eagle’s Medium (DMEM) with 10% FBS. GM9503 primary skin fibroblasts (from a healthy 10 year old male donor) were obtained from the Coriell Institute (Camden, NJ). They were cultured in Minimum Essential Medium (MEM) with 10% FBS. All cell lines were maintained at 37°C and 5% CO2 atmosphere.

### Cell growth studies

To assess the effects of ACSVL3 inhibitors on the growth of U87MG cells, 5000 U87MG or U87-KO cells were seeded into wells of 6-well plates containing DMEM + 10% FBS (U87MG cell culture media) on day 0. On day 1, media was aspirated and replaced with cell culture media containing either CB5 or CB16.2 (1:1000 dilution of a 1000x stock solution in DMSO) or equivalent DMSO only (control). The final concentration(s) of drug were as indicated in each figure legend. Triplicate wells of each condition were harvested via gentle trypsinization and counted using a hemocytometer on days 3, 6, 9, 12, and 15. Cell media and drug/DMSO were refreshed every 3 days.

To assess reversion of growth inhibition in U87MG cells previously treated with CB5, the culture medium was replaced on day 9 with medium containing DMSO alone. Culture was continued throughout the remainder of the experiment. Triplicate wells were harvested via trypsin and counted by hemocytometer on days 9, 12, 15, 18, 21, and 23. Cell media and drug/DMSO were refreshed every 3 days.

### Cell morphology

5000 U87MG or U87-KO U87 cells were seeded onto 3.5 cm plates on day 0. On day 1, half were treated with 3 μM CB5 as described earlier and half were treated with equivalent DMSO. To assess changes in morphology induced by CB5, cells were imaged with using light microscopy with an Axiovert 100 TV microscope (Carl Zeiss) on days 3, 6, 9, and 12. Cell media and drug/DMSO were refreshed every 3 days.

Human primary fibroblasts (GM9503) were seeded into wells in a 12-well plate at approximately 80% confluency. The next day, duplicate wells of cells were treated with the following concentrations of CB5: 1 μM, 3 μM, 6.5 μM, 10 μM, and DMSO only. Cells were allowed to grow for 5 days and then imaged with an EVOS XL Core Imaging System (Thermo Fisher Scientific). Media and drug were refreshed on the 3rd day.

### Fatty acid β-oxidation assay

U87MG cells were seeded into 6-well plates and allowed to grow to near confluence. For each fatty acid, duplicate blank wells (media only, no cells) were set up to serve as extraction controls.

Approximately 1.5 hour before the assay, media was removed and replaced with 540 μl serum-free DMEM containing 2.2 mM carnitine and 3 μM CB5 (3 mM stock in DMSO diluted 1:1000 in media) or equivalent DMSO for control and blank wells. Replicate wells were set up to determine protein concentration using the Pierce 660nm reagent (Thermo Fisher).

The assay measures the release of labeled water-soluble metabolites from a [1-14C]fatty acid resulting from one round of β-oxidation. Stock solutions containing 100 μM [1-14C]fatty acid (20 μM radiolabeled fatty acid plus 80 μM unlabeled fatty acid) in benzene were prepared, dried under a stream of nitrogen and solubilized in an equal volume of 10 mg/ml α-cyclodextrin (Millipore Sigma) in 10 mM Tris pH 8.0. 60 μl of each solubilized fatty acid was added to duplicate wells containing U87MG cells treated with CB5, U87MG cells treated with DMSO, and blank, cell-free wells. Plates were manually shaken gently to mix and incubated at 37°C for 2 hours. The incubation was stopped by adding 120 μl of ice-cold 18% HClO4 to each well. Water-soluble products of β-oxidation were separated from unreacted fatty acids using the method of Folch [Folch et al., 1957]. Radioactivity in the upper, aqueous phase of each extraction was quantitated by liquid scintillation counting. Activity expressed as nmol fatty acid degraded/2h/mg protein was calculated and normalized to the activity measured with C16:0 in untreated U87MG cells.

### Immunofluorescence and western blotting

Cells were seeded into culture plates containing glass coverslips. The next day, media containing 3 μM CB5 or equivalent DMSO was added. Cells were treated for 9 days. Media and drug/DMSO were refreshed every 3 days. Coverslips were removed from culture plates and GFAP immunofluorescence measured as previously described [Jia et al., 2007]. The primary α-GFAP antibody (Dako) was diluted 1:5000 in 0.01% BSA in PBS and the secondary antibody was Cy3-conjugated goat-α-rabbit (Jackson ImmunoResearch) diluted 1:150 in 0.01% BSA in PBS. Cells were visualized by fluorescence microscopy using an Axio Imager M2 with ApoTome attachment (Carl Zeiss).

Expression of GFAP in CB5-treated and untreated CB5 was also assessed by western blotting as previously described [Laemmli, 1970; Jia et al., 2007]. The primary antibody used was α-GFAP (1:500 dilution in 10% nonfat dry milk solution; Dako). β-Actin was also detected as a loading control using α-β-actin antibody (1:800 dilution in milk; Santa Cruz Biotechnology). The secondary antibody used was goat α-rabbit IgG-HRP (1:8000 dilution in milk solution; Santa Cruz Biotechnology).

### Scratch-wound healing assay

Cells were seeded into 6-well plates and allowed to grow to about 80% confluence. A plastic 1000 μl pipet tip was used to scratch the cell layer in each well. Scratches were imaged immediately after the scratch, 24 hours post-scratch and 48 hours post-scratch to assess cell growth over the scratch wound (healing). Imaging was performed with an EVOS XL Core Imaging System (Thermo Fisher Scientific).

### Animals and their care

All animal studies were approved by the Johns Hopkins University School of Medicine Institutional Animal Care and Use Committee (IACUC) in accordance with the guidelines and regulations described in the NIH Guide for the Care and Use of Laboratory Animal. Female NOD/SCID mice were obtained from the Johns Hopkins University Sidney Kimmel Comprehensive Cancer Center Office of Research Services Animal Resources and were housed in the Cancer Center’s animal facility throughout the experiments. Mice were maintained in a pathogen-free environment at a constant temperature (22°C) on a 12-hour light/dark cycle with ad libitum access to food and water.

### Toxicity of CB5 in mice

CB5 was dissolved at a concentration of 20 mg/mL into DMSO. CB5/DMSO was diluted into sterile PBS and sonicated in a water-bath type sonicator (Branson) for 15 minutes at 30°C to form a stable suspension. (Sonication was performed before injections each day). Concentrations used were 10, 20, 40 and 80 μl of CB5/DMSO per mL for the doses of 2, 4, 8, and 16 mg/kg/day, respectively. Two 8-week old female NOD/SCID mice each received increasing doses of CB5 via a 200 μl intraperitoneal injection once daily. They received 2 mg/kg for 2 days, 4 mg/kg for 2 days, 8 mg/kg for 6 days, and 16 mg/kg for 2 days. Animals were observed for 30 minutes following each injection to determine any acute side effects of CB5.

Toxicity was also assessed using Kolliphor HS 15 (12-Hydroxy-octadecanoicacid polymer with α-hydro-ω-hydroxypoly(oxy-1,2-ethanediyl); Millipore Sigma) as a vehicle. A 30% solution (w/v) was made of Kolliphor HS 15 in PBS. The solution was sterilized by filtration. CB5 was dissolved directly into the Kolliphor solution. Two 12-week old female NOD/SCID mice each received the following doses of CB5 in a daily 200 μl injection: 2, 4, 8, 16, 32, and 64 mg/kg. Each dose was given for two days, and the final 64 mg/kg dose was given for 5 days. Animals were observed for several minutes after injections to determine any acute side effects.

### U87MG xenografts

Near-confluent U87MG cells were harvested via trypsin and counted. 4 x 10^6^ cells in 50 μl sterile PBS were injected subcutaneously into both hind flanks of six 8-week old female NOD/SCID mice. Tumors were allowed to grow and measured every 2-3 day using calipers. Tumor volume was approximated using the following formula: volume = (length x width^2^)/2. Once average tumor size was approximately 100 mm^3^, mice were injected daily with 2 mg/kg CB5/DMSO in PBS prepared as described for toxicity studies. CB5 was given to 3 mice via intraperitoneal injection once daily. The other 3 mice received equivalent volume of DMSO in PBS (vehicle). Mice receiving vehicle were sacrificed on Day 22, when average tumor volume exceeded 1 cm^3^. Mice receiving CB5 were sacrificed on Day 25 when a large outlier tumor volume exceeded 1 cm^3^. Tumors were excised and weighed after sacrifice.

### Mayo 22 xenografts

Mayo 22 cells were derived from a primary glioblastoma and are maintained by serial passage as subcutaneous xenografts [Carlson and Sarkaria, 2011]. To implant new xenograft tumors, 500 mg was measured out from the previous generation’s subcutaneous xenograft. This tumor section was finely chopped, suspended in 5 mL of sterile PBS, and passed 6 times through a 16-gauge needle using a 10 mL syringe. The suspension was centrifuged, aspirated, and resuspended in 2 mL of PBS. 100 μl of this suspension was implanted by subcutaneous injection into both hind flanks of 8-week old female NOD/SCID mice. On day 9, when average tumor was approximately 100 mm3, half of the mice received daily injection of 8 mg/kg CB5/DMSO in PBS prepared as described for toxicity studies; the remaining mice received vehicle alone. Mice were sacrificed on Day 23 once some tumor volumes exceeded 1 cm^3^. Tumors were excised and weighed.

### Statistical analysis

Unless otherwise specified, all results are presented as mean and SD of triplicate observations. Where appropriate, a two-tailed Student’s t-test was used to calculate statistical significance.

## RESULTS

### CB5 slows the growth rate of U87 cells in monolayer culture

U87MG cells exhibit malignant growth properties including rapid growth, lack of contact inhibition when grown in monolayer culture and adherence-independent growth in soft agar suspension [Pei et al., 2009; Kolar et al., 2021]. They also form rapidly growing xenografts when injected subcutaneously or orthotopically implanted into nude mice [Pei et al., 2009; Kolar et al., 2021].

Depletion of ACSVL3 by either siRNA-mediated knockdown or genetic knockout reverts U87 cells to a more normal (i.e. less malignant) growth phenotype [Pei et al., 2009; Kolar et al., 2021]. We therefore hypothesized that U87MG cells treated with a small drug-like molecule that inhibits ACSVL3 enzyme activity should have a growth phenotype more like that exhibited by knockdown or KO cells. As no ACSVL3 inhibitors have been reported, we sought to identify one or more candidates. We recently reported that CB5, also known as grassofermata, significantly inhibited ACSVL3 enzyme activity in U87MG cells [Clay et al., 2025].

Due to its dramatic effects on acyl-CoA synthetase activity, the effect of CB5 on growth of glioma cells in culture was characterized. First, 10 µM CB5 was added to media of U87MG cells in culture. This appeared to be toxic to the cells. Many detached over the course of several days or developed a rounded, sick-looking morphology. When the CB5 concentration was reduced to 3µM, the drug had no negative effect on cell adherence or mortality. However, at this concentration CB5 significantly slowed the growth rate of U87MG cells (Figure 1A). U87MG cells grown in 3 μM CB5 grew significantly slower than not only U87MG cells treated with DMSO but also U87-KO cells. 3 μM CB5 appeared to be somewhat toxic to U87-KO cells, as the number of cells began to decrease slightly after 9 days of treatment.

**Figure 1:**
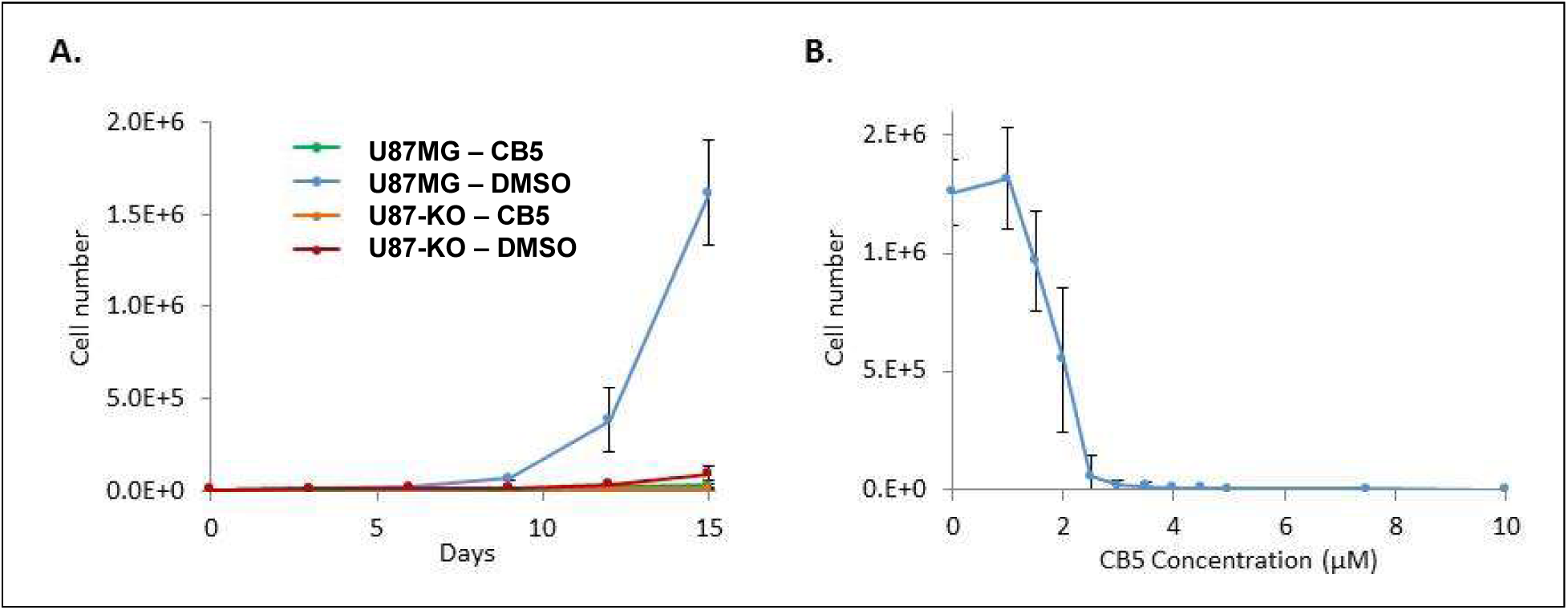
CB5 inhibits cell proliferation in both U87MG and U87-KO cells. A. U87MG and U87-KO cells were seeded into wells (5000 cells/well) of 6-well plates on day 0. On day 1, medium containing either CB5 in DMSO (final concentration 3 µM) or DMSO alone was added. Cells from triplicate wells were counted every 3 days for 15 days. CB5 dramatically decreased the growth rate of both cell lines when measured at either day 12 (p<0.05) or day 15 (p<0.001). Similarly, CB5 also lowered the rate of U87-KO cell proliferations on both day 12 (p<0.01) and day 15 (p<0.05). Notably, the rate of proliferation of CB5-treated U87MG cells was also significantly lower than the rate of untreated U87-KO cells (p<0.05 for both day 12 and day 15). B. U87MG cells were treated with increasing concentrations of CB5 for 12 days, after which cells were counted. The therapeutic range of CB5 on U87MG cells *in vitro* is narrow; 1 µM or less had no effect on proliferation rate, while 10 µM or more completely suppressed cell proliferation. 3 µM CB5, the dose used for other *in vitro* experiments using U87MG cells, is near the lower point of inflection of the S-shaped dose response curve. Results are presented as the mean number of cells from triplicate wells ± SD.

To assess dose dependence, U87MG cells were treated for 12 days with increasing concentrations of CB5. Growth was found to be inhibited by CB5 in a dose-dependent manner (Figure 1B). The effective range of CB5 appeared to be very narrow; while 1 μM CB5 had no effect on growth, 10 μM was toxic to U87MG cells in culture.

### CB5 affects the morphology of U87MG cells in culture

We previously reported that U87-KO cells had altered morphology compared to U87MG cells [Kolar et al., 2021]. The ACSVL3-deficient KO cells appeared to be larger and flatter than U87MG cells, and take up more surface area (Figure 2). Therefore, we asked whether treating U87MG cells with CB5 would produce similar changes. When U87MG cells were incubated with 3μM CB5, morphological changes were indeed seen (Figure 2). However, CB5-induced morphological changes were different from the changes observed in U87-KO cells. After several days of growth in the presence of CB5, U87MG cells appeared to be larger and grew longer projections compared to vehicle-treated cells (Figure 2). These changes are consistent with the hypothesis that CB5 may induce differentiation of U87MG cells.

**Figure 2.**
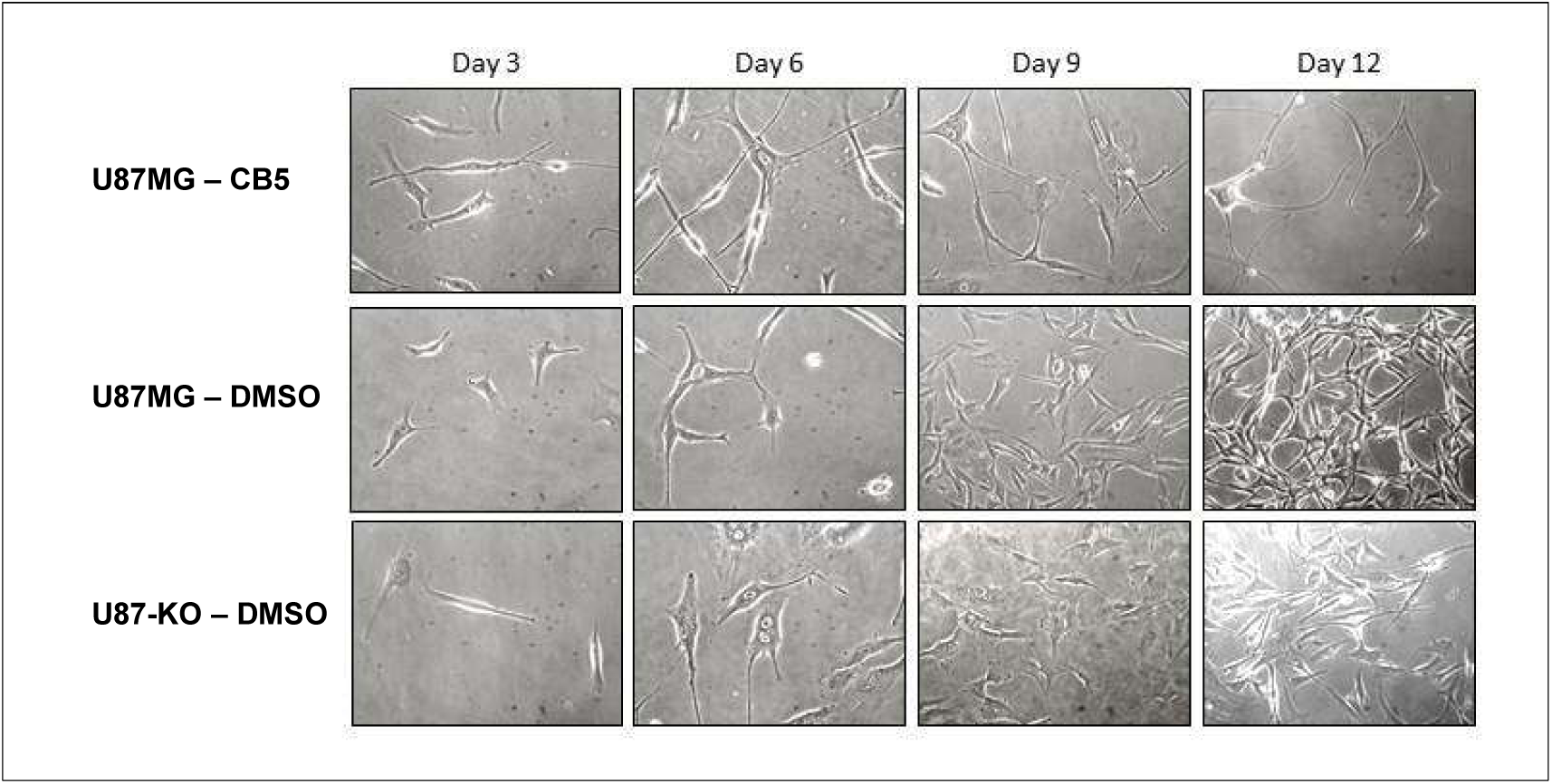
CB5 treatment does not alter the morphology of U87MG or U87-KO cells. U87MG cells were treated with 3 µM CB5 for 12 days and were then examined using differential interference contrast (DIC) microscopy. CB5 treatment induced a unique morphology in U87MG cells. The cells were larger and grew long projections. The morphology of U87-KO cells is shown for comparison. ACSVL3-deficient U87-KO cells also have a morphology distinct from U87MG cell morphology, and it is also dissimilar to the morphology induced by CB5. Representative micrographs from triplicate plates are shown.

### CB5 induces differentiation of U87MG cells

To determine whether CB5 induces differentiation of U87MG cells, expression of the astrocyte-specific marker glial fibrillary acidic protein (GFAP) was assessed by immunofluorescence and western blot. CB5-treated cells showed higher levels of GFAP in U87MG cells treated for 9 days with 3μM CB5 compared to cells treated with DMSO (Figure 3). While RNAi knock-down of ACSVL3 led to an increase of GFAP in several low passage primary GBM neurosphere cell lines directly derived from patient samples [Sun et al., 2014], the U87-KO line did not exhibit this phenotype by immunofluorescence (Figure 3).

**Figure 3:**
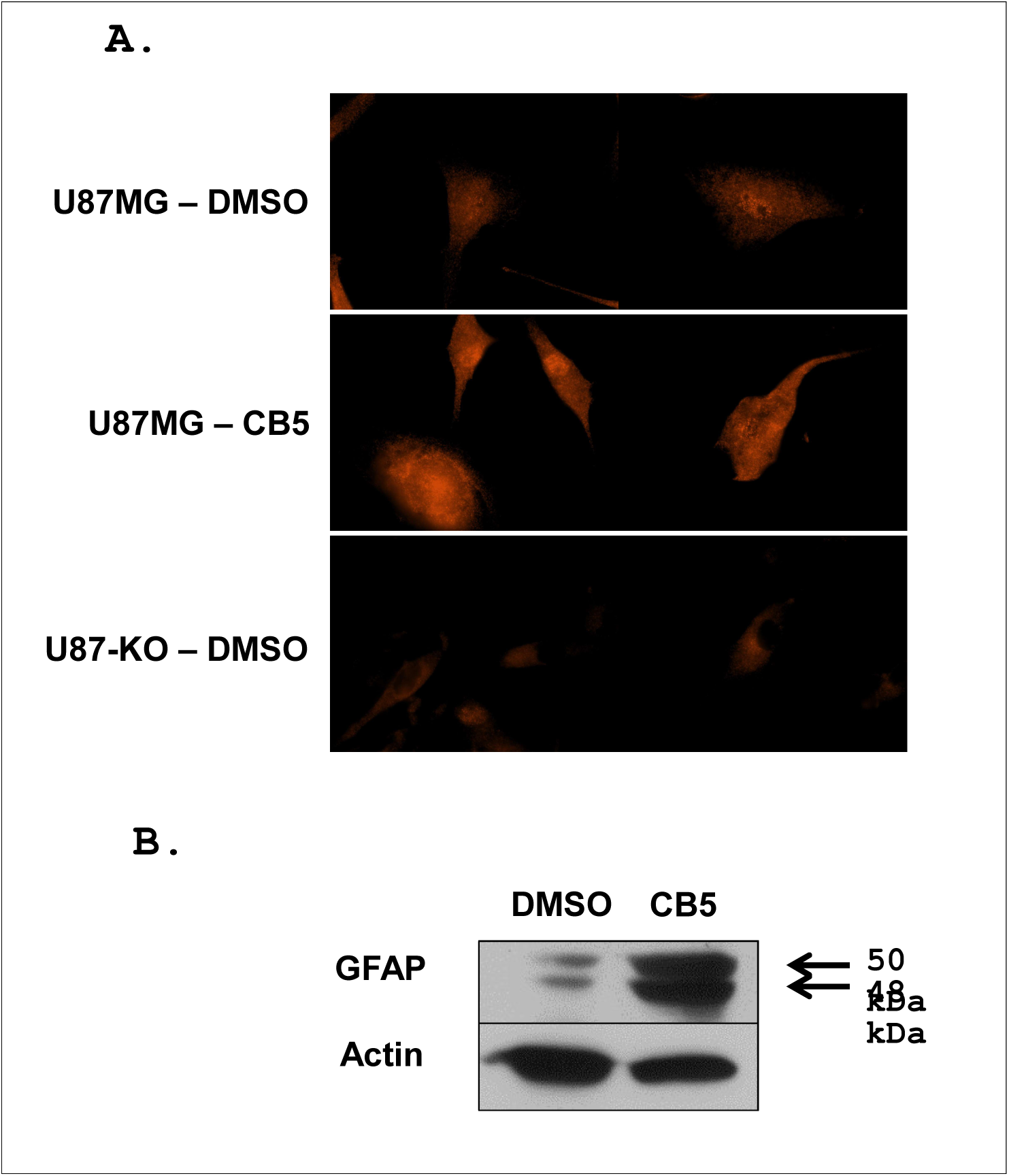
CB5 induces differentiation in U87MG cells. Expression of the astrocyte-specific protein GFAP was evaluated in U87MG cells treated with 3 µM CB5 or DMSO for 9 days, and U87-KO cells. A. Indirect immunofluorescence revealed that GFAP, a marker of differentiation, was more abundant in CB5-treated U87MG cells than either DMSO-treated U87MG cells or DMSO-treated U87-KO cells. B. Western blotting confirmed that 50 kDa GFAP and a proteolytic fragment of ∼48 kDa were much more abundant in CB5-treated U87MG cells than DMSO-treated U87MG cells.

### CB5 did not affect cell growth or morphology of primary human cells

A desirable property of an anti-neoplastic drug is that normal tissue not be adversely affected at a therapeutic concentration. To test the effect of CB5 on a non-cancerous cell line, primary human fibroblasts in culture were treated with increasing concentrations of CB5 (1-10 µM) for 5 days. At all concentrations tested, CB5 had no obvious effect on cell growth or morphology (Figure 4), even at concentrations that were toxic to U87MG cells. Notably, human skin fibroblasts do not express detectable ACSVL3 protein (The Human Protein Atlas; proteinatlas.org).

**Figure 4:**
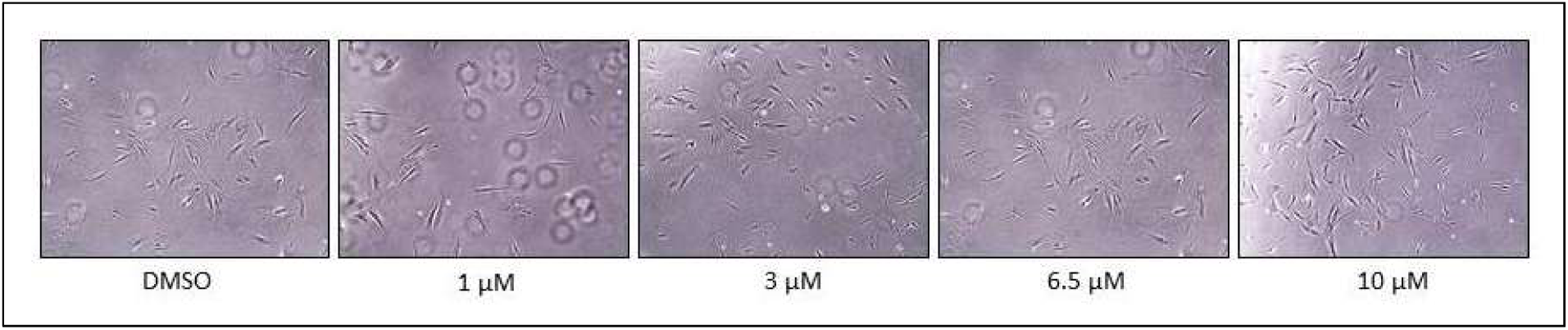
CB5 is not toxic to human fibroblasts. Normal human skin fibroblasts were treated with varying concentrations of CB5 and observed for 5 days. None of the concentrations tested, including 10 µM CB5 (which was toxic to U87MG cells), had an effect on cell number or morphology of the fibroblasts.

### The effect of CB5 on U87MG cell growth is reversible

To determine whether 3 μM CB5 was cytotoxic to U87MG cells, we grew the cells for 9 days in the presence of this drug. Media containing CB5 was removed from half of the groups of cells and replaced with media containing the DMSO vehicle only. The U87MG cells from which CB5 was removed continued to grow slowly for several days, but the rate eventually increased to an exponential rate (as seen in U87MG cells treated with DMSO alone) around Day 21, or 12 days after CB5 was removed (Figure 5). Thus, growth inhibition by CB5 appeared to be reversible and not cytotoxic to U87MG cells at the concentration tested.

**Figure 5:**
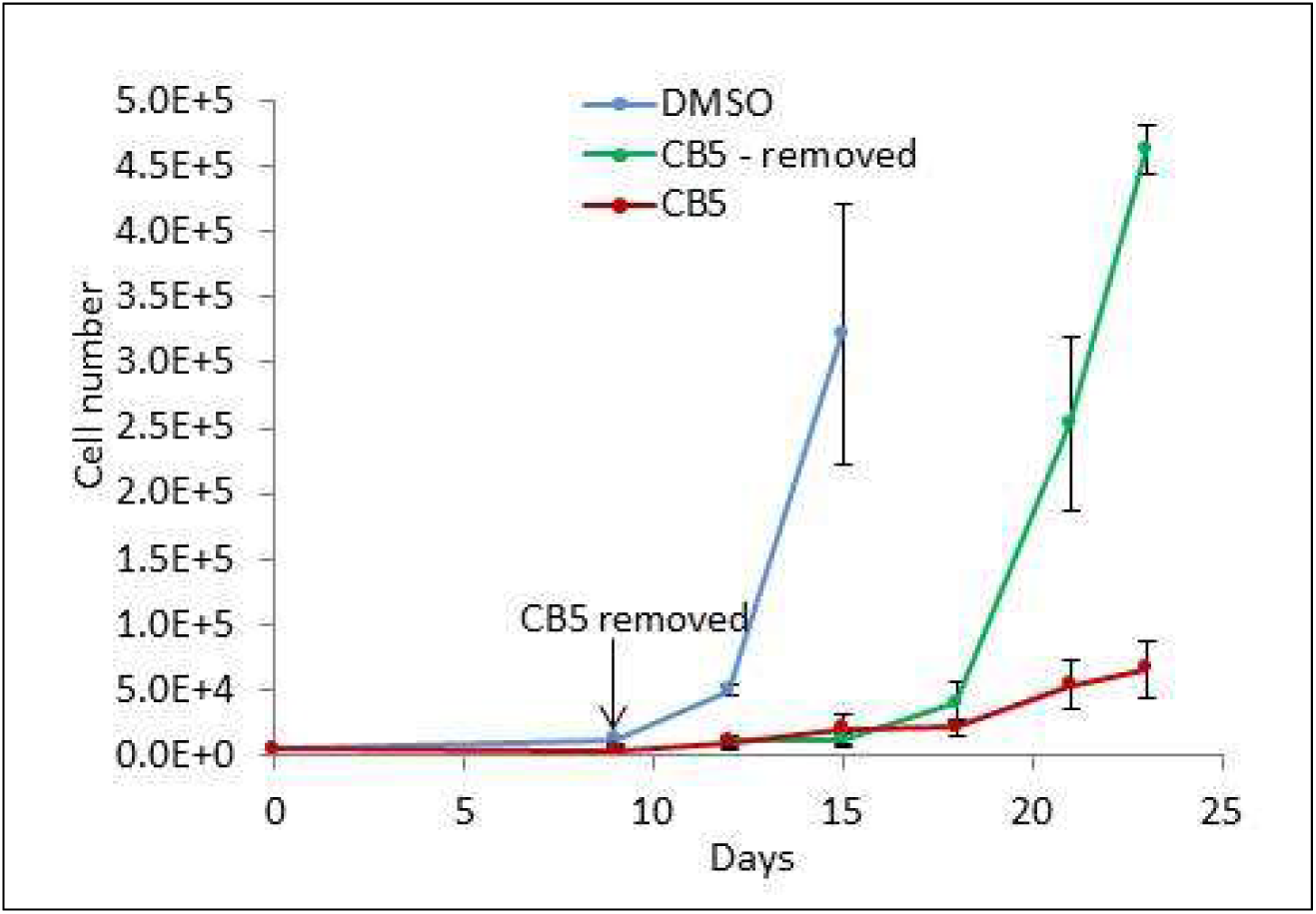
The depressed growth rate of CB5-treated U87MG cells reverts to that of untreated U87MG cells following drug removal. U87MG cells were treated with DMSO or 3 µM CB5 in DMSO for 9 days. At that time, CB5 was removed from half of the treated cells and replaced with medium containing DMSO only. Cells were counted every 3 days. Several days after removal, U87MG cells that had been previously treated with CB5 resumed the untreated growth rate. Results are presented as the mean number of cells from triplicate wells ± SD. The decreased growth rate of U87MG cells treated continuously with CB5 compared to untreated U87MG cells was significant on days 12 (p<0.001) and 15 (p=0.017), recapitulating the findings in Figure 1. The accelerated growth rate of U87MG cells following removal of CB5 was significantly higher than that of continuously treated cells (day 21, p<0.05; day 23, p=0.013).

### CB5 decreased invasiveness of U87MG cells as measured by the scratch-wound assay

Invasiveness is a primary characteristic of malignant cells. Thus, a desirable property of an anti-tumor drug is to reduce invasiveness of the tumor cells. The scratch-wound assay was used to compare the invasiveness of U87MG cells treated with 3 μM CB5 to that of untreated U87MG and U87-KO cells. For each condition, a layer of cells growing in culture was scratched with a pipette tip, and the ability of the cells to close the wound was assessed. After 48 hours, the scratch in the cells treated with CB5 remained visible, while the scratch in the DMSO-treated U87MG cells had closed (Figure 6A).

**Figure 6.**
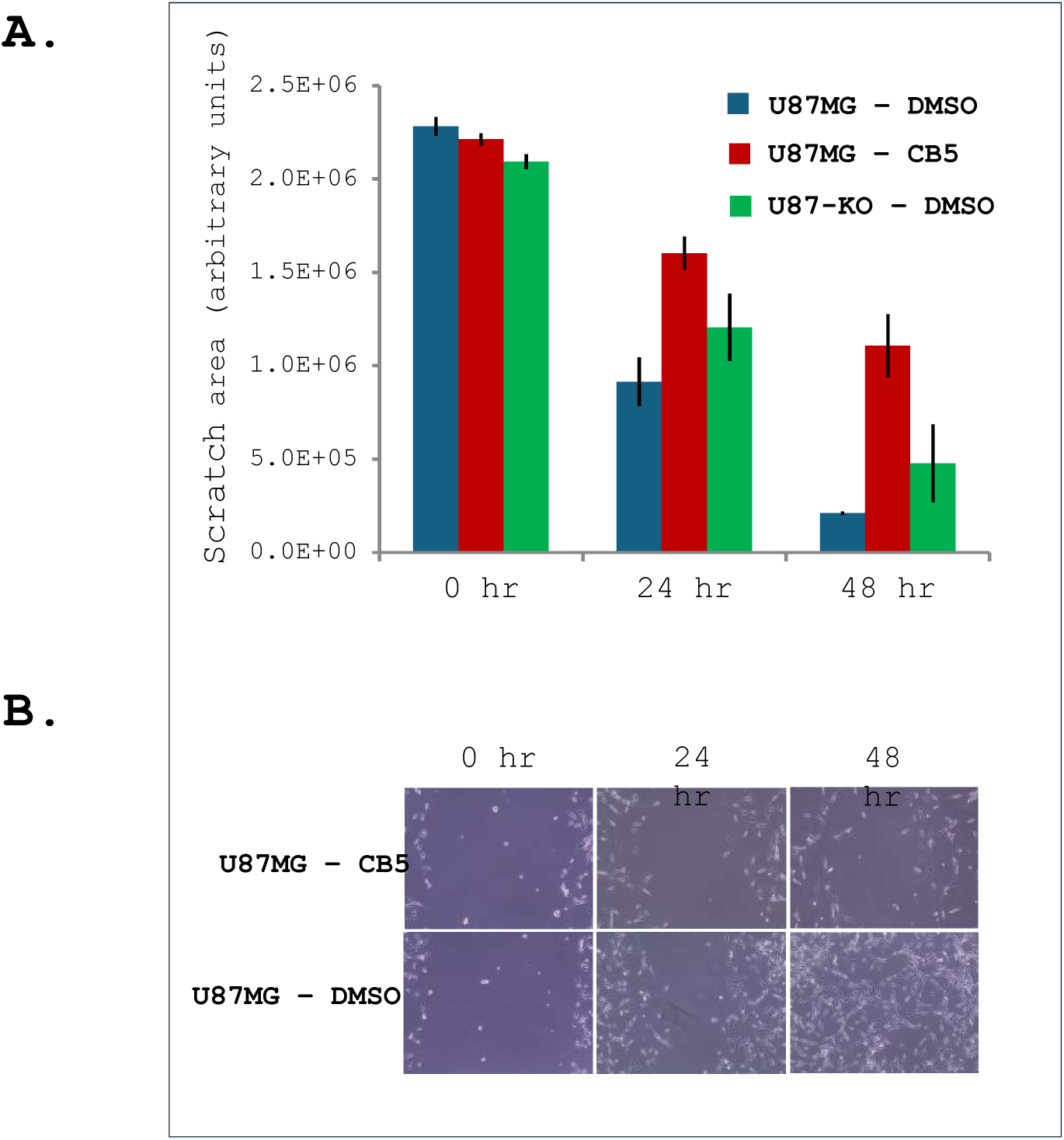
Invasiveness of U87MG cells is mitigated by treatment with CB5. The scratch-wound assay was used to assess invasion of U87MG cells treated with either 3 µM CB5 or DMSO. U87-KO cells treated with DMSO only were also compared. Quantification of the scratch sizes in each condition (A) and images of representative scratches in CB5- and DMSO-treated U87MG cells (B) are shown. In CB5-treated U87MG cells, the scratch closed more slowly than in either DMSO-treated U87MG cells (p=0.002 at 24 h and p=0.0014 at 48 h) or U87-KO cells (not significant). Scratch sizes were quantified using ImageJ software [Schindelin et al., 2015].

Quantification of the scratch size reveals that DMSO-treated U87MG cells closed the scratch most rapidly, followed by DMSO treated U87-KO cells. CB5-treated U87MG cells closed the scratch the slowest (Figure 6B). While this assay does not demonstrate whether cell migration or cell division is a bigger factor in closing the scratch, it does suggest that CB5-treated U87MG cells are less invasive than both U87MG cells and U87-KO cells treated with DMSO alone.

### CB5 acutely decreases β-oxidation of long-chain fatty acids in U87MG cells

Once activated to its CoA derivative by an ACS such as ACSVL3, a fatty acid can participate in many catabolic and anabolic pathways. Degradation of long-chain and very long-chain FA by β-oxidation in mitochondria and peroxisomes, respectively is one such pathway. To determine whether CB5 affects the β-oxidation of long- and very long-chain fatty acids, U87MG cells were incubated with 3 μM CB5 or DMSO vehicle alone for 1.5 h, followed by the addition of radiolabeled fatty acids. After an additional 2 h incubation, water-soluble products of β-oxidation were quantified. Long-chain FA tested were palmitic acid (C16:0), stearic acid (C18:0) and oleic acid (C18:1); VLCFA tested were behenic acid (C22:0) and lignoceric acid (C24:0). All results were normalized to the rate of palmitic acid β-oxidation in DMSO-treated U87MG cells.

The rate of palmitic, stearic, and oleic acid β-oxidation was reduced when CB5 was present (Figure 7). β-Oxidation of behenic acid appeared to be only slightly reduced by CB5. While lignoceric acid degradation trended toward reduction, the signal-to-noise ratio was high preventing interpretation of this result with confidence. These results indicate that in addition to its longer-term effects on growth and morphology, CB5 acutely affects FA metabolism in U87MG cells.

**Figure 7.**
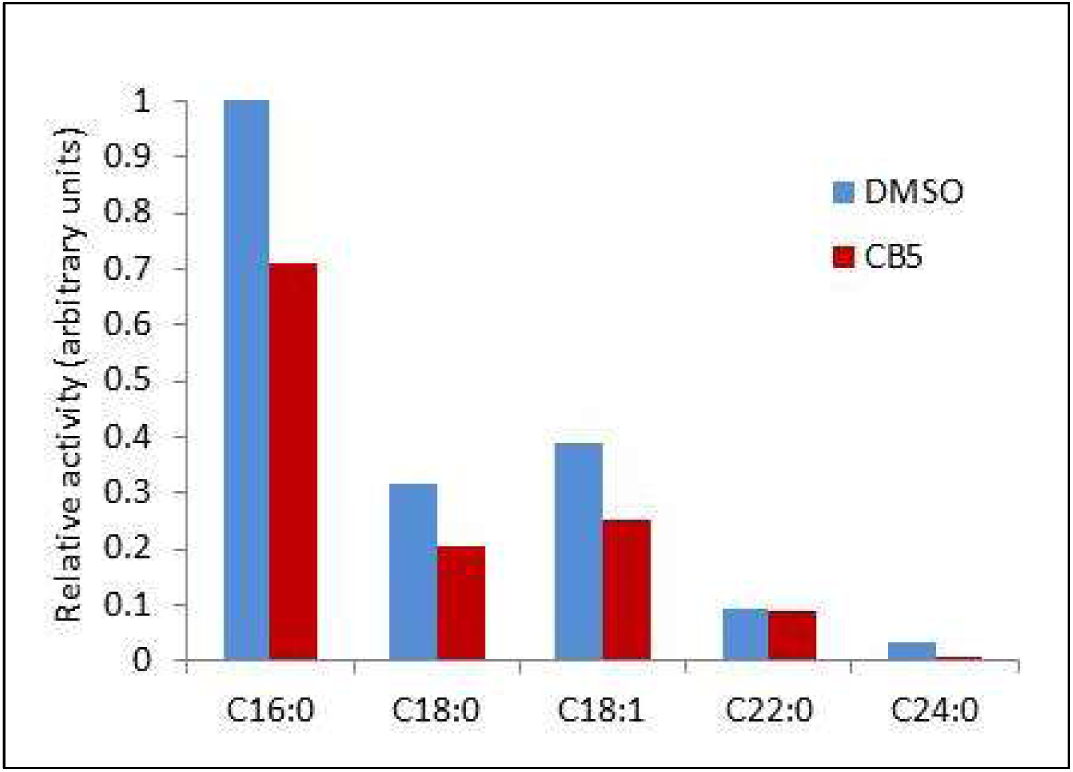
CB5 acutely decreases β-oxidation of long- and very long-chain fatty acids in U87MG cells. U87MG cells were treated with either 3 µM CB5 or vehicle DMSO alone for 1.5 hr prior to the addition of [1-^14^C]-labeled fatty acids. Release of labeled water-soluble β-oxidation products over a 2 hr. period was quantitated. All results were normalized to the rate of C16:0 oxidation in DMSO-treated U87MG cells. CB5 decreased the rate of β-oxidation of each FA tested, particularly the long-chain FAs C16:0, C18:0, and C18:1.

### CB5 is well tolerated by mice

Another compound that inhibits growth of U87MG cells that we had considered for further study is CB16.2, also known as lipofermata [Sandoval et al., 2010; Clay et al., 2025]. However, CB16.2 was toxic to mice in doses larger than 2 mg/kg when given by intraperitoneal injection; mice became immobile for several minutes after dosing (C. DiRusso, University of Nebraska-Lincoln, personal communication). To assess whether CB5 is toxic to mice under similar conditions, nude mice were treated with increasing concentrations of CB5 administered by daily intraperitoneal injection. CB5 was dissolved in DMSO, which was then diluted into PBS to form a stable suspension. NOD/SCID mice were treated with 2 mg/kg/day for 2 days, then 4 mg/kg/day for 2 days, 8 mg/kg/day for 6 days, and finally 16 mg/kg/day for 2 days. CB5 seemed to be better tolerated that CB16.2. At no point did the mice exhibit any obvious side effects, including the CNS side effects reported with larger doses of CB16.2.

To assess whether CB5 was more bioavailable to mice in solution than in suspension, we dissolved CB5 in Kolliphor HS 15. NOD/SCID mice were treated sequentially for two days each with 2, 4, 8, 16, and 32 mg/kg CB5 administered by intraperitoneal injection. At all concentrations tested, CB5 was entirely solubilized. Finally, the mice were treated for 5 days with the highest dose tested, 64 mg/kg/day. This concentration did not go fully into solution but did form a stable suspension. The mice did not exhibit obvious side effects from any dose. Although one desired outcome of this study was to determine a toxic dose of CB5, concentrations higher than 64 mg/kg did not form stable suspensions.

### CB5 slows tumor growth in mice bearing U87MG xenografts but not in Mayo-22 xenografts

To determine whether CB5 treatment affects growth of U87MG or Mayo-22 cell subcutaneous xenografts, NOD/SCID mice were injected subcutaneously on both hind flanks with either U87MG cells (Figure 8A and 8B) or Mayo-22 cells (Figure 8C and 8D). When tumors had grown to an average volume of 100 mm^3^, half of the mice received daily injections of CB5 (2 mg/kg/day) while the other half got daily injections of an equivalent volume of DMSO. Tumor size was measured every 2-3 days. After 7 days of treatment (15 days post-xenograft implantation), U87MG tumors in the mice treated with DMSO began to grow more rapidly than those treated with CB5 (Figure 8A). After 12 days of treatment (20 days post-implantation), tumors in the mice treated with CB5 began, on average, to decrease in size. The tumor sizes exhibited by the CB5-treated mice (except for one outlier, where z = 2.0 for final tumor weight) then stagnated. DMSO-treated mice were sacrificed on post-injection day 22 and CB5-treated mice on day 25. Despite the extra 3 days of tumor growth, weights of harvested tumors from CB5-treated mice were on average much lower than tumors from DMSO-treated mice (Figure 8B). Of the 6 total tumors from mice treated with CB5, one was considered to be an outlier as it grew at a similar rate to the tumors from the DMSO-treated mice. Paradoxically, the tumor on the mouse’s other flank responded similarly to the other tumors from CB5-treated mice.

**Figure 8.**
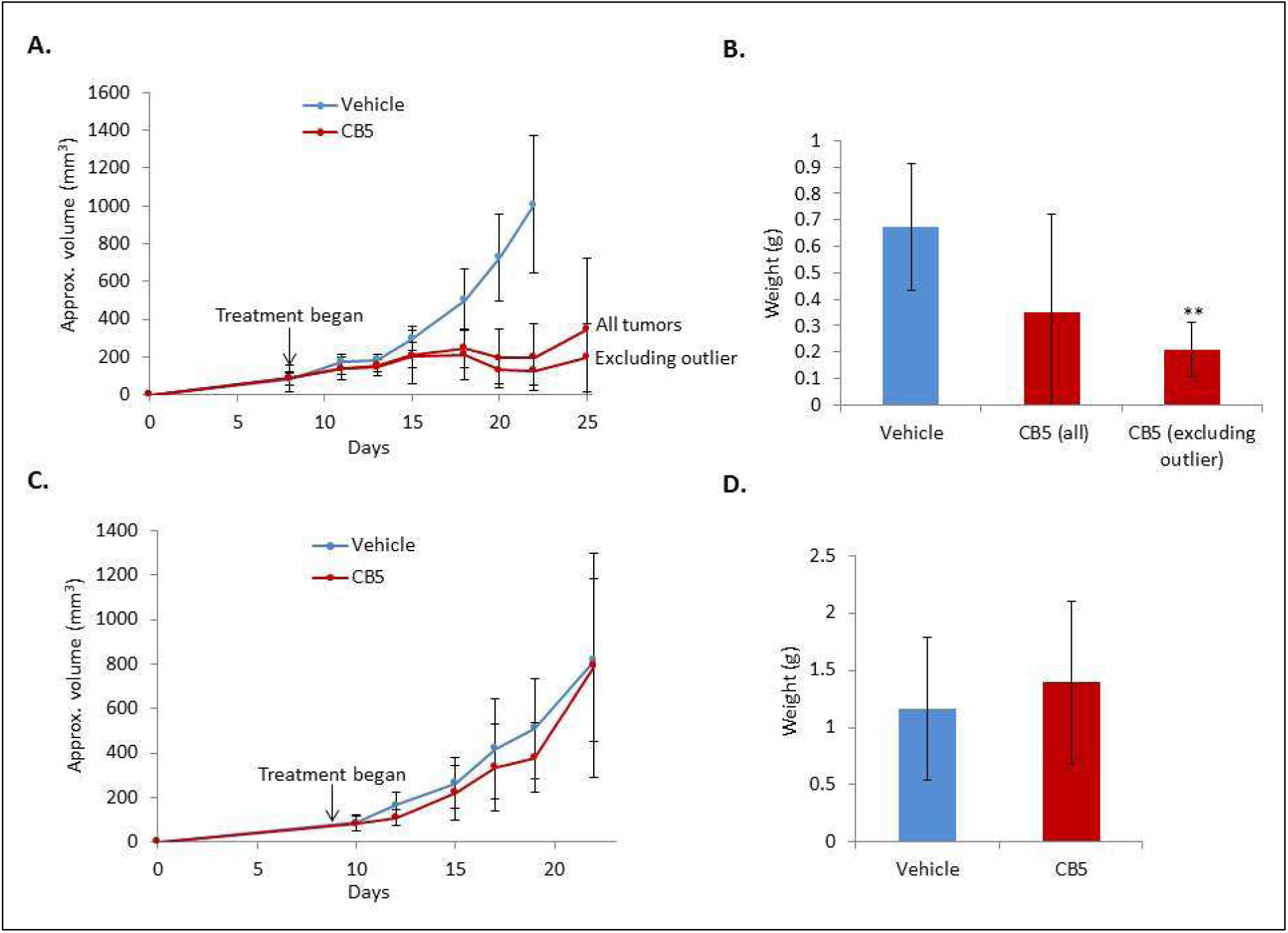
CB5 decreases growth of subcutaneous xenograft tumors from U87MG cells but not Mayo 22 cells. A. Mice bearing subcutaneous xenograft tumors formed from U87MG cells were treated with 2 mg/kg/day CB5 in DMSO or the equivalent volume of DMSO beginning 8 days after tumor implantation. Tumor size was approximated every 2-3 days by measuring length and width with calipers. B. Vehicle-treated mice were sacrificed 22 days after xenograft implantation, and CB5-treated mice were sacrificed 25 days after injection (n=6 for each condition). Tumors were excised and weighed. If one outlier was excluded (see text), the tumors from CB5-treated mice grew slower (p<0.01) and even decreased in volume over the course of treatment. Final tumor weight after sacrifice is shown. C. Mice bearing tumors formed from Mayo-22 cells were treated with 8 mg/kg/day CB5 beginning 9 days after tumor implantation. Approximate tumor size is shown. Mice were sacrificed 22 days after tumor implantation. There was no significant difference in approximate tumor size at any time point or final tumor weight after sacrifice (D).

Xenografts arising from GBM cells maintained in vivo as subcutaneous xenografts, such as Mayo-22 cells, are thought to represent better the situation in human tumors [Patrizii et al., 2018]. Therefore, we injected another cohort of NOD/SCID mice subcutaneously with Mayo 22 cells and tumors were allowed to grow to approximately 100 mm^3^ (Figure 8C and 8D). Half of the mice then received daily injections of CB5, this time at a higher dose of 8 mg/kg, while the other half received an equivalent volume of DMSO. Tumors were measured every 2-3 days. Mayo-22 tumor growth was significantly more variable than that observed with U87MG xenografts within both the CB5-treated mice and the group receiving DMSO. Thus, there was no statistical difference between DMSO-treated and CB5-treated mice with respect to either estimated tumor volume or final tumor weight (Figure 8C and 8D).

## DISCUSSION

Acyl-CoA synthetases play a central role in lipid metabolism by catalyzing the thioesterification of CoA to a FA, thereby “activating” it for subsequent metabolism [Watkins, 1997]. While investigating the properties of very long-chain ACSs (ACSVLs) that are capable of activating FAs containing 22 or more carbons, we made the unexpected observation that ACSVL3 was overexpressed in glioma cells including GBM [Pei et al., 2009]. Further studies in U87MG and other glioma cell lines in which ACSVL3 expression was either knocked down by RNA interference or knocked out by genetic deletion suggested that many malignant properties of glioma cells were dependent on ACSVL3 expression [Pei et al., 2009; Kolar et al., 2021]. We therefore asked whether an inhibitor of ACSVL3s enzymatic activity would act in a similar anti-malignant fashion, reducing its enzyme activity while maintaining the abnormally high protein expression observed in glioma cells. Since no ACSVL3 inhibitors were known, we sought to identify one or more candidates. Such a tool would be useful not only for its therapeutic potential, but also for investigating the biochemistry and physiologic role(s) of ACSVL3 in glioma metabolism.

The U87-KO cell line has been very useful in characterizing the role that ACSVL3 plays in lipid metabolism, carbohydrate metabolism and cell cycle in glioma [Kolar et al., 2021; Kolar et al., 2023; Yang et al., 2024], and halting ACSVL3 activity with an inhibitor should in theory produce the same phenotypes exhibited in the U87-KO line. While most results presented here support this hypothesis, not all do. For example, CB5-treated U87MG cells show an increase in GFAP expression vs. DMSO-treated U87MG cells by immunofluorescence, while U87-KO cells did not show an increase (Figure 3). Some phenotypic differences between U87-KO cells and CB5-treated U87MG cells are likely attributable to compensatory changes that arise in the KO line to combat the loss of ACSVL3; proteomic analysis revealed that many proteins are upregulated or downregulated in the U87-KO line compared to WT U87s [Kolar et al., 2021], and while some of these changes are likely induced by CB5, using an inhibitor allows investigation of the mechanism(s) by which ACSVL3 affects the glioma phenotype without the need to consider the effects of compensatory changes that arise over time.

In studies reported previously [Clay et al., 2025] we identified several compounds that inhibited ACSVL3, but none were specific for this enzyme. For therapeutic use, a highly specific inhibitor would be most desirable. However, for initial studies assessing the anti-neoplastic potential of an inhibitor, a high degree of specificity is not critical. A high-throughput screen of chemical and drug libraries conducted by Black and associates had identified inhibitors of a related enzyme, ACSVL1 [Sandoval et al., 2010; Li et al., 2007], and we tested several of these compounds for their ability to inhibit ACSVL3 [Clay et al., 2025]. One of these compounds, CB5 (designated “grassofermata” by the Black laboratory [Saini et al., 2015]), significantly inhibited the ACS activity of ACSVL3 when assayed in U87MG cells with radiolabeled stearic acid (C18:0), a preferred ACSVL3 substrate. In addition, the fractional inhibition of stearoyl-CoA synthesis by CB5 was smaller in U87-KO cells, as would be expected for cells lacking ACSVL3 [Clay et al., 2025]. Based on these observations, a more in-depth analysis of CB5’s anti-malignant properties seemed warranted. Therefore, we investigated growth rate, scratch wound healing, and tumorigenic potential of U87MG cells cultured with CB5.

Similar to the effect of ACSVL3 depletion by knockdown or KO, the growth rate of U87MG cells in culture was significantly decreased when CB5 was present in the culture medium (Figure 1A). While untreated U87MG cells began to show significant growth beginning at 7-8 days in culture, cells treated with 3 µM CB5 showed no observable growth even after 15 days in culture. CB5 did not kill the treated U87MG cells, however, as removal of the compound restored cell growth after several days in culture (Figure 5).

Interestingly, CB5 had a narrow range of effectiveness. 1 μM CB5 had no effect on U87MG cell proliferation, while 10 μM was toxic to these cells. The effect of CB5 on proliferation formed an S-curve when cell number was plotted against CB5 concentration (Figure 1B). A concentration of 3 μM, which was used for most of the experiments performed on U87MG cells in culture, was found to be at the lower point of inflection on this S-curve. This suggests that it may be an ideal does to use on cells in culture, balancing toxicity with effectiveness. Interestingly, the toxicity seen in U87MG cells treated with high doses of CB5 was not observed in non-cancerous cells. When normal primary human skin fibroblasts were treated for 5 days with CB5 at doses up to 10 μM, no changes in either cell number or morphology were found (Figure 4). This suggested that the drug may target tumor cells without affecting healthy cells, which is ideal for cancer therapeutic agents.

CB5 treatment also altered the morphology of U87MG cells. When treated with 3 µM CB5 for 12 days, U87-KO cells appeared to be larger, and grew long projections (Figure 2). These morphological changes caused us to question whether CB5 was inducing differentiation in these cells. Thus, we looked for changes in expression of the differentiation marker, GFAP. Indeed, we found increased GFAP in CB5-treated U87MG cells compared to DMSO-treated cells by both immunofluorescence and western blotting.

As noted above, restoration of wild-type U87MG cell growth after removal of CB5 from the culture medium took several days. This raised the question as to how this occurs, given that CB5 induces differentiation of U87 cells. One hypothesis is that increased proliferation stems from a smaller, sub-population of CB5-treated cells and thus explains why it takes several days to revert to the untreated rate. After several days of treatment with CB5, there were a few wild-type U87MG cells that did not have the altered morphology induced by CB5, and when GFAP expression was evaluated via immunofluorescence, occasional cells were found that exhibited less GFAP. This hypothesis could also explain why an outlier xenograft U87 tumor (Figure 8) did not respond to CB5 treatment. Future experiments will aim to characterize this heterogeneity in treated cells.

Invasiveness of tumor cells is a marker of malignancy that can be assessed in culture by scratch-wound healing. While U87MG cells readily migrated into a scratch wound made with a plastic pipet tip, treatment with CB5 significantly decreased this rate, consistent with a less malignant phenotype (Figure 6). Overall, studies presented here are consistent with our hypothesis that similar to knocking out ACSVL3 (i.e. the U87-KO cell line), inhibiting the enzyme activity of ACSVL3 with CB5 creates a more normal, and thus less malignant, cellular phenotype.

The Black laboratory that initially identified ACSVL1 (FATP2) inhibitors used decreased transport of the fluorescent FA C1-BODIPY-C12 rather than decreased enzyme activity as the screening criterion [Sandoval et al., 2010]. When they pre-treated CaCo-2 cells for 1 hour with inhibitor and then assayed ACSL1 enzyme activity with [^14^C] oleate as substrate, only CB2 at high concentration was inhibitory [Sandoval et al., 2010]. They concluded that the inhibitors targeted domains important for FA transport, but not ACS enzyme activity [Sandoval et al., 2010]. In contrast, our initial screening tool was the inhibition of ACS enzyme activity [Clay et al., 2025]. Inhibiting ACSVL3 by CB5 should result in decreased rates of downstream metabolic processes such as FA β-oxidation. This makes sense, since fatty acids must be converted to acyl-CoA prior to β-oxidation. Because rapidly proliferating tumor cells need to generate significant amounts of energy, impairing β-oxidation may be one of the ways that CB5 slows proliferation of U87 cells. Indeed, treatment of U87MG cells with 3 µM CB5 decreased the rate of FA β-oxidation by CB5 (Figure 7). However, a direct effect of CB5 on one or more β-oxidation enzymes cannot be ruled out. Further experimentation will be conducted to sort this out.

U87MG cells form subcutaneous tumors in mice. We had previously shown that KO of ACSVL3 in these cells reduced the number of xenografts that formed and the sizes of these tumors [Kolar et al., 2021] and we wondered whether CB5 might have similar effects. However, side effects and tolerability are important considerations for patients undergoing treatment with chemotherapeutic agents [Shrestha et al., 2019; Islam et al., 2019]. We therefore tested the effects of increasing concentrations of CB5 on w.t. mice. CB5 was well tolerated by mice and did not appear to cause any physical or behavioral side effects at the IP doses tested, up to 64 mg/kg/day.

When we assessed the effect of 2 mg/kg/day CB5 on weights of U87MG subcutaneous xenografts (Figures 8A and 8B), we observed a reduction similar to that seen in xenografts produced from ACSVL3-deficient U87-KO cells [Pei et al., 2009; Kolar et al., 2021]. However, the decrease was statistically significant only when one totally non-responding tumor was removed from the analysis. It is not clear why there was no effect of CB5 on this tumor, as a xenograft growing in the opposite flank of the same animal responded to CB5.

We also found that tumors generated from another xenograft-producing glioma line, Mayo-22, did not respond to CB5 treatment even at a higher dose of 8 mg/kg/day (Figure 8C and 8D). The different responses may be explained by physiological differences between the two types of tumors, such as vascularization. Another possibility is that CB5 is inhibiting tumor growth in U87 xenografts not by inhibition of ACSVL3 but by targeting another pathway. CB5 is also a known inhibitor of ADP-ribosylation factor 6 (Arf6), a GTP-binding protein that regulates endocytic recycling and cytoskeleton remodeling (World Patent No. 2015183989). It is possible that CB5 could be inhibiting Arf6 in U87-based xenografts but not in Mayo 22-based xenografts. Future directions include evaluating the level of Arf6 in both U87 and Mayo 22 cells.

Another concern with CB5 treatment is the short half-life of the drug. CB5 has a half-life of 151.7 minutes in blood when given intravenously (C. DiRusso, University of Nebraska-Lincoln, personal communication). While IP injection likely leads to a longer half-life, better results may be achieved with multiple injections per day or a continuously-injecting, implanted micro-pump. Using a vehicle such as Kolliphor HS-15 (Millipore Sigma), which was used to characterize CB16.2 in mice, may affect the potency of the drug as well. Future experiments will test different delivery systems and higher concentrations of CB5 on Mayo 22-based xenografts.

It is not yet known if CB5 will cross the blood-brain barrier, which would be ideal for treating gliomas. Future experiments will be needed to determine whether treating mice bearing intracranial xenografts with is effective at reducing tumor growth rates.

## ACKNOWLEGEMENTS

The authors thank Drs. Paul Black and Concetta DiRusso (Dept. of Biochemistry, Univ. of Nebraska, Lincoln NE) for providing several ACSVL1 inhibitors used in this study and for sharing unpublished data. This work was supported by NIH grant NS062043.

## Notes

### Competing Interest Statement

The authors have declared no competing interest.

